# How the immune mousetrap works: structural evidence for the immunomodulatory action of a peptide from influenza NS1 protein

**DOI:** 10.1101/2023.12.25.573289

**Authors:** Yana Zabrodskaya, Vladimir Tsvetkov, Anna-Polina Shurygina, Kirill Vasyliev, Aram Shaldzhyan, Andrey Gorshkov, Alexander Kuklin, Natalya Fedorova, Vladimir Egorov

**Affiliations:** Smorodintsev Research Institute of Influenza, Russian Ministry of Health, 15/17 Ulitsa Prof. Popova, St. Petersburg 197376, Russia; Institute of Biomedical Systems and Biotechnology, Peter the Great Saint Petersburg Polytechnic University, 29 Ulitsa Polytechnicheskaya, St. Petersburg 194064, Russia; Federal Research and Clinical Center for Physical Chemical Medicine, 1a Ulitsa Malaya Pirogovskaya, Moscow 119435, Russia; Institute of Biodesign and Complex System Modeling, Sechenov First Moscow State Medical University, 119146 Moscow, Russia; International Intergovernmental Organization Joint Institute for Nuclear Research, 6 Ulitsa Joliot-Curie, Dubna 141980, Russia; Moscow Institute of Physics and Technology (State University), 9 Institutskiy pereulok, 141701 Dolgoprudny, Moscow Region, Russia; Petersburg Nuclear Physics Institute named by B. P. Konstantinov of National Research Center, Kurchatov Institute, 1 mkr. Orlova roshcha, Gatchina 188300, Russia; Institute of Experimental Medicine, 12 Ulitsa Akademika Pavlova, St. Petersburg 197376, Russia

**Keywords:** T-cell receptor, NS1, influenza A virus, fibrillogenesis, immunosuppression, conformational transition

## Abstract

One of the critical stages of the T-cell immune response is the dimerization of the intramembrane domains of T-cell receptors (TCR). Structural similarities between the immunosuppressive domains of viral proteins and the transmembrane domains of TCR have led several authors to hypothesize the mechanism of immune response suppression by highly pathogenic viruses: viral proteins embed themselves in the membrane and act on the intramembrane domain of the TCRalpha subunit, hindering its functional oligomerization. It has also been suggested that this mechanism is used by influenza A virus in NS1-mediated immunosuppression. We have shown that the peptide corresponding to the primary structure of the potential immunosuppressive domain of NS1 protein (G51) can reduce concanavalin A-induced proliferation of PBMC cells, as well as in vitro, G51 can affect the oligomerization of the core peptide corresponding to the intramembrane domain of TCR, using AFM and small-angle neutron scattering.

The results obtained using *in cellulo* and *in vitro* model systems suggest the presence of functional interaction between the NS1 fragment and the intramembrane domain of the TCR alpha subunit. We have proposed a possible scheme for such interaction obtained by computer modeling.

This suggests the existence of another NS1-mediated mechanism of immunosuppression in influenza.

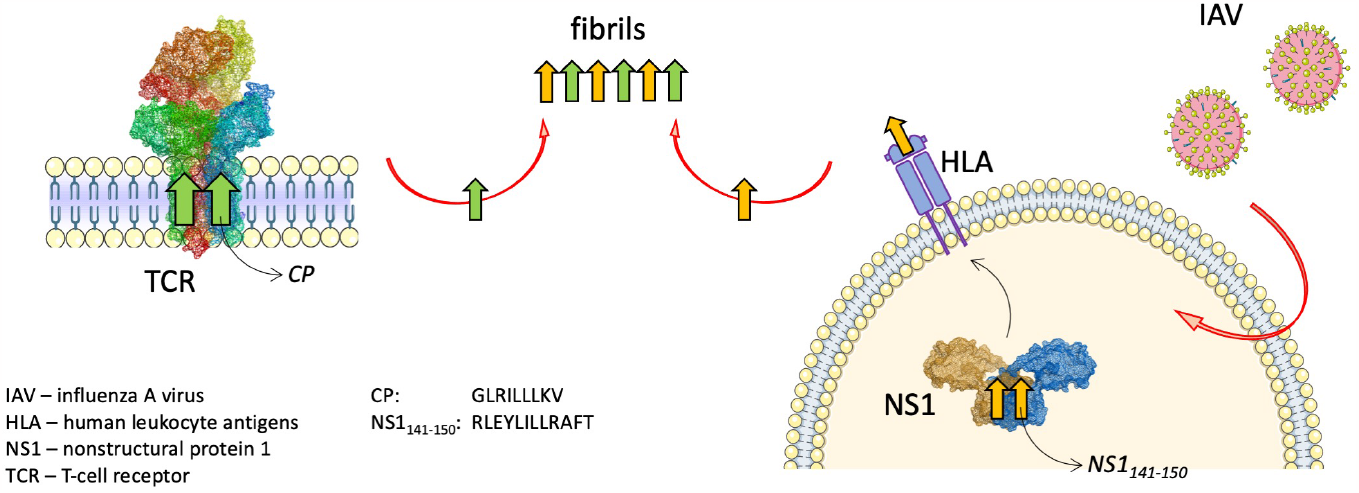

## 1. Introduction

The NS1 protein of influenza A virus is one of the products of the viral RNA NS segment. Depending on the strain, the length of the NS1 polypeptide chain varies from 215 to 237 amino acid residues. The protein consists of two domains: an RNA-binding domain (RBD, residues 1-70) and an effector domain (ED, residues after 80), connected by a short linker sequence. Among the known strains of influenza A virus, the C-terminal domain of NS1 is the most variable [1]. In addition to RNA, NS1 protein has a wide range of cellular partners [2,3].

The functions of NS1 protein can be conditionally divided into three groups: 1) suppression of the interferon and antiviral response of the cell; 2) inhibition of processing and export of cellular mRNAs; 3) promotion of translation of viral mRNAs, regulation of the activity of the viral polymerase [4].

NS1 inhibits the antiviral protection of infected cells by preventing the activation of interferon-inducing proteins and by inhibiting the action of effector proteins [5–7], or by blocking splicing and export of cellular mRNAs from the nucleus to the cytoplasm [8–11]. Currently, it has been established that NS1 interacts with proteins which can modulate gene expression, including through chromatin remodeling [12–14].

Viruses with remote RNA fragments encoding NS1 protein are incapable of replicating in cells that produce interferon [15]. Amino acid substitutions in NS1 of different strains of influenza A virus correlate with their pathogenicity [16]. The currently known mechanisms for suppressing the immune response by the NS1 protein are extensive, but do not fully describe its role in the development of viral infection. NS1 is localized in the nucleus of cells at early stages of infection, and in the cytoplasm at later stages [17]. Recent studies have shown the presence of NS1 protein in the virion, but the number of molecules it comprises averages 4.2 ± 3.2 molecules per 1 molecule of polymerase complex (strain A/WSN/33, MDBK cells) [15]. This suggests a nonspecific, random incorporation of this protein into the virion in low quantities during its formation when it is highly concentrated in the cytoplasm. Thus, secreted extracellular NS1 (free or in virions) can affect various components of the immune system. Moreover, it should be noted that fragments of NS1 are represented on the surface of cells by the major histocompatibility complex [18].

One critical stage of T-cell immune response is the dimerization of intramembrane domains of T-cell receptors (TCR), which initiates the phosphorylation of tyrosine kinases of the cytoplasmic domains of immunoreceptors. This provides intracellular signal transmission and cell proliferation necessary for the formation of an immune response. In [19] the structural similarity of immunosuppressive domains of viral proteins and transmembrane domains of TCRs is described, and a hypothesis about the mechanism of immune response suppression by highly pathogenic viruses is raised. According to this hypothesis, viral proteins are embedded in the membrane and act on the intramembrane domain of the alpha subunit of the T-cell receptor, preventing its oligomerization necessary for phosphorylation and signal transduction [20]. It has been shown that even relatively short polypeptides, such as the core peptide CP (**GLRILLLKV, CP**), corresponding in primary structure to the intramembrane domain of the alpha subunit of the T-cell receptor, are capable of immunosuppressive action by such a mechanism. In [1] it was shown that such a peptide is capable of modulating T-cell response by interacting with components of the receptor complex within the membrane, replacing one of the full-size receptor subunits. The monograph [20] suggests that a similar mechanism of action on T-cell receptors can be involved in NS1-mediated immunosuppression by the influenza A virus, and that an NS1 fragment homologous to CP may also prevent activation of T-cell immunity by blocking antigen-induced receptor oligomerization and consequently, triggering the signaling cascade leading to T-cell division.

It should be noted that CP forms aggregates and beta-structured oligomers even at high concentrations and in the presence of negatively charged membranes *in vitro* [21]. Two-dimensional NMR experiments (TOCSY and NOESY) have shown the formation of beta-structures in aqueous solutions at low pH. Upon incorporation into the membrane as a monomer, the peptide undergoes a conformational transition [22]. Also, using surface plasmon resonance, it was shown that the CP peptide can interact with artificial analogues of T-cell membranes and with isolated T-cell membranes [23].

However, the sequence of the potential immunosuppressive domain of NS1, given in [20] corresponds to the core region of amyloid-like fibrils formed by the NS1 protein *in vitro* [24]. The presence of homology in the sequences of CP and the potential immunosuppressive domain, together with the ability of both peptides to form beta-structured oligomers, and in the case of NS1 fragment - amyloid-like fibrils, provides a basis for suggesting that such peptides may influence each other’s conformation by a prion-like mechanism. Recent studies have shown the presence of a peptide domain among the peptides represented by the major histocompatibility complex on the surface of infected cells [18], indicating that direct contact between such a peptide and T-cell receptors is possible during antigen recognition by T-cells *in vivo*. Thus, the propensity of the potential immunosuppressive domain to form amyloid-like fibrils, together with the propensity of the TCRalpha intramembrane domain to form beta-structured oligomers and the homology in the amino acid sequences of TCRalpha and NS1 ISD domains may lead to the induction of conformational transitions of many TCR receptors on the surface of T-cells.

It should be noted that *NS* gene has been detected in extracellular vesicles secreted by influenza virus-infected cells, but its function remains unclear [25]. However, the role of extracellular vesicles that carry proteins with altered conformations in the spread of conformational diseases is widely discussed [26].

In summary, it was shown that the transmembrane domain of the T-cell receptor (CP peptide) is capable of interacting with the entire receptor within the membrane. At the same time, in previous studies exploring the potential immunosuppressive effects of viruses due to the homology of their fragments with CP, the mechanism of cleavage of a peptide capable of being integrated into the membrane was not considered. The crucial factor that enabled the use of the peptide model in this work was the discovery of the homologous CP fragment of influenza NS1 among the peptides presented by MHC on the cell surface [18].

To demonstrate the fundamental possibility of the operation of such an immunosuppressive mechanism during influenza, in this work we investigated the effect of a peptide corresponding to the primary structure of the potential NS1 ISD on the induced proliferation of PBMC cells and studied the influence of the corresponding peptide on the oligomerization of the CP peptide *in vitro*.

## 2. Materials and methods

### 2.1 Peptides

Peptides GLRILLLKV (CP) and RLEYLILLRAFT (G51) were synthesized by LLC “Verta”. Sigma-Aldrich (USA) reagents were used.

### 2.2 Preparation of peptide solutions

#### CP and G51 peptides

50 μl of 100% TFE (trifluoroethanol, Sigma) was added to dry CP or G51 peptide, and then after complete dissolution of the peptide, 450 μl of water was added. The final concentration of peptides was 1 mM. Peptides were incubated for 24 hours at 55ºC with constant shaking in an Eppendorf thermomixer (USA) at 300 rpm.

#### Peptide mixture

25 μl of 100% TFE was added to dry CP or G51 peptide, then the solutions were mixed, after that 450 μl of water was added. The peptide mixture was incubated for 24 hours at 55ºC with constant shaking in an Eppendorf thermomixer (USA) at 300 rpm.

### 2.3 Carboxyfluorescein diacetate succinimidyl ester (CFSE) dilution assay

Human peripheral blood mononuclear cells (PBMCs) were isolated by Ficoll-Hypaque density gradient centrifugation from heparinized peripheral blood (buffy coat) under sterile conditions and cryopreserved. Prior to CFSE (Sigma) staining and subsequent stimulation, PBMCs were thawed and left to rest overnight seeded into the 48 well culture plates (2*10^6^ cells per well) in full culture medium: RPMI 1640 medium supplemented with 10% heat inactivated FBS and 1% penicillin-streptomycin mixture (all reagents supplied by Gibco).

For CFSE staining cells were harvested into centrifugation tube, counted, pelleted and stained in 1 ml of 5μM CFSE solution for 5 min at room temperature and then 5 min on ice. The staining was quenched by adding 5 volumes of ice-cold full culture medium. The cells were pelleted, washed and plated in 48-well culture plates at a concentration of 1 × 10^6^ cells/well. CFSE-labelled cells were incubated for 1 h at 37°C, 5% CO_2_ with serial dilutions of G51 peptide or 1% DMSO solution in full culture medium and then stimulated with concanavalin A (ConA, Sigma) in a final concentration 2.5 μg/ml for 5 days. As a control of proliferation inhibition doxorubicin was used at the concentration 1 μg/ml. Addition of doxorubicin was made on the third day of incubation. Following incubation cells were stained with a dead cell marker (LIVE/DEAD Zombie Aqua stain; Biolegend) and anti-CD4-APC/Cy7 (BD), anti-CD3-PE/Cy7 (Beckman Coulter) and anti-CD8-PerCPCy5.5 (BD) monoclonal antibodies, fixed and acquired on a BD Canto II flow cytometer. Data analysis was performed using FCSExpress (De Novo software).

### 2.4 Transmission electron microscopy (TEM)

A sample (10 μl) was applied to a formvar-coated copper mesh. After 1 minute of incubation, the sample was collected and washed twice with water. For negative staining, the sample-containing mesh was incubated for one minute with 20 μl of phosphotungstic acid. Excess acid was removed with a piece of filter paper. Electron microphotographs were obtained using a JEOL JEM 1011 electron microscope (Japan).

### 2.5 Atomic force microscopy (AFM)

The samples were 10-fold diluted and put on freshly cleaved mica (SPI Supplies) at a volume of 10 μl. After 1 minute incubation at room temperature, the mica surface was washed three times with water to remove salt. The sample topography measurements were performed in semi-contact mode on an atomic force microscope (NT-MDT– Smena B) using the NSG03 probe (NT-MDT, Russia). The images were analyzed using Gwyddion software [27].

### 2.6 Congo red assay (CR)

Congo red dye (Sigma Aldrich, USA) at a concentration of 50 μM in PBS was mixed with the 10 μM peptide. As a control, a similar sample was used in which a buffer was added instead of the peptide. Absorption spectra were recorded on an Avantes Ava Spec 2048 instrument (Avantes, The Netherlands). Congo red dye spectra demonstrate characteristic right-shift when mixing with samples containing amyloid-like fibrils.

### 2.7 Thioflavin T (ThT)

To obtain fluorescence spectra, a solution of Thioflavin T (25 μM) was added to PBS, followed by addition of 5 nmol of the peptide solution (or a mixture of peptides, 5 nmol each). Fluorescence was recorded on a Hitachi F-7000 FL spectrofluorometer (Japan) with excitation at 440 nm (slit width 5 nm) and fluorescence detection at wavelengths between 450-700 nm (slit width 5 nm).

### 2.8 Chromatography

For chromatographic separation of peptides by gel filtration, a YMC-Pack Diol 60 column (YMC, Japan) with dimensions of 500×8 mm and a granule size of 5 μm was used. The mobile phase consisted of 10% TFE (Sigma, USA) in PBS, with a flow rate of 1 ml/min. A sample of 10 μl was applied. Detection was performed at a wavelength of 220 nm.

### 2.9 Small-angle neutron scattering (SANS)

To obtain SANS spectra, solutions of CP, G51 peptides, and their mixtures were prepared at a concentration of 2 mg/ml in 5% DMSO/PBS/D_2_O. Peptides were first dissolved in 100% DMSO, mixed, if necessary, then the required amount of 10x PBS/D_2_O and D_2_O was added. Samples were incubated at 55°C for one hour on an Eppendorf orbital shaker. Spectra were obtained using a YuMO spectrometer located on the fourth channel of the high-throughput reactor IBR-2 (Frank Neutron Physics Laboratory, Joint Institute for Nuclear Research, Dubna, Russia). Measurements were carried out in the standard geometry described in [28,29]. Scattering was registered with two detectors simultaneously in the range of q = 0.006-0.3 Å^-1^. Data preprocessing was carried out according to the scheme described in [30].

All spectra were visualized using Origin2015 software and processed using the GNOM program (ATSAS package) [31] to calculate of the distance distribution function of the cross-section P(R) assuming monodisperse system of rod-like particles. The function *P*(*R*) = *γ*_*c*_(*R*) · *R* is evaluated, where *γ*_*c*_(*R*) is the characteristic function of thickness. The visualization of scattering objects based on SANS data was performed according to the ATSAS manual using DAMMIN [32] as was described earlier [33].

### 2.10 Mass spectrometry

The samples were mixed with DHB matrix (2,5-dihydroxybenzoic acid, Bruker, Germany), put on Ground steel plate and dried. The spectra were obtained at MALDI-TOF/TOF mass spectrometer UltrafleXtreme (Bruker) in reflective and linear positive ions registration modes.

### 2.11 Computer modelling

The following strategy was used to construct fibril models. At the first stage, by means of the docking procedure using the Molsoft package ICM-Pro 3.9-2 [34], the mechanism of interaction between the peptides CP and G51 during the formation of complexes CP-CP, G51-G51 and CP-G51 was studied and the most probable geometry of the location of the contacting peptides was determined. Then, the two-peptide complex obtained at the first stage was docked with its copy. Similarly, the four-monomer complex was docked with its copy, and so on, until the complex was obtained from the required number of peptides. During the docking procedure, force field ECEPP [35] and the biased probability Monte Carlo minimization procedure were used.

The stability of the resulting fibril models was verified by the molecular dynamics method using the Amber 20 package [36]. An influence of the solvent simulated with application model of water molecules TIP3P. The simulation performed by using periodical boundary conditions and rectangular box. The buffer between the models and the periodic box wall was at least 15 Å. To mimic a neutral physiological environment, an ionic environment was added in two steps. First, chlorine ions were added to neutralize the charge due to the fibril structures. Then chlorine and sodium ions were added in amounts to simulate a neutral physiological environment. The parameters needed for interatomic energy calculation were taken from the force fields ff14SBonlysc [37]. At the beginning of computing the investigated systems minimized by two steps. At the first stage location of the solvent molecules optimized by using 1000 steps (500 steps of steepest descent followed by 500 steps of conjugate gradient), at that mobility of all solute atoms was restrained with a force constant of 500 kcal*mol^-1^*Å^-2^. At second stage the optimization realized without any restriction using 2500 steps (1000 steps of steepest decent, 1500 steps of conjugate gradient). Then gradual heating to 300 K during 20 ps was performed. To avoid wild fluctuations for investigated systems in this stage weak harmonic restrains were used with a force constant of 10 kcal*mol^-1^*Å^-2^ for all atoms being not part of solvent. SHAKE algorithm was applied to constrain bonds to hydrogen atoms, that allowed to use 2 fs step. Scaling of nonbonded 1–4 van der Waals and electrostatic interactions were performed by the standard amber values. The cutoff distance for non-bonded interactions was equal to 10 Å and the long-range electrostatics calculated using the particle mesh Ewald method. The MD simulations in production phase were carried out using constant temperature (T = 300 K) and constant pressure (p = 1 atm) over 100 ns. To control the temperature Langevin thermostat was used with the collision frequency of 1 ps^-1^. Berendsen barostat was used to control the pressure. The free energy was calculated as the sum of the electrostatic energies (E_q_), Van der Waals energies (E_VDW_), energy of solvation and deformation energy of valence bonds, valence and dihedral angles (U). The energy of solvation was calculated as the sum of the polar and nonpolar contributions. The polar contribution (E_GB_) was computed using the Generalized Born (GB) method and the algorithm developed by Onufriev et al. for calculating the effective Born radii [38]. The non-polar contribution to the solvation energy (E_surf_), which includes solute-solvent van der Waals interactions and the free energy of cavity formation in solvent, was estimated from a solvent-accessible surface area (SASA). The plots of geometrical parameters and energy of interaction vs. time were smoothed using the moving average method (span = 5).

### 2.12 Graphical abstract

For graphical abstract NS1 structure 4OPH [39] and TCR structure 7FJD [40] were used; an Influenza-virus icon, an emptycell-membrane-halfcircle icon, a membrane-2d-yellowblue icon by Servier https://smart.servier.com/ are licensed under CC-BY 3.0 Unported https://creativecommons.org/licenses/by/3.0/.

## 3. Results

### 3.1 Effect of the G51 peptide on the proliferation of T-lymphocytes

A peptide containing a potential ISD from influenza virus NS1 was synthesized and its effect on induced T-cell proliferation was studied.

The effect of G51 peptide on T-cell proliferation was studied in carboxyfluorescein diacetate succinimidyl ester (CFSE) dilution assay using cryopreserved human peripheral blood mononuclear cells (PBMCs). The optimal dosage of concanavalin A to induce proliferation in linear range was determined in preliminary experiment and amounted to 2.5 mg/ml. Prior stimulation CFSE-labeled PBMCs were incubated with serial dilutions of G51 for 1h then ConA was added and cells were left for 5 days to proliferate. Analysis of CD4+ and CD8+ lymphocytes proliferation was performed by flow cytometry.

It was shown that G51 peptide inhibits the ConA-induced multiplication of T lymphocytes in dose dependent manner (Fig.1).

Thus, we have shown that the peptide corresponding to the primary structure of the potential ISD from the NS1 protein of the influenza virus can cause dose-dependent inhibition of PBMC cell proliferation induced by concanavalin A. To determine the possible mechanism of such inhibition, we studied the possibility of interaction between the studied peptide and the core fragment of the TCRalpha receptor in vitro in a membrane-mimicking environment.

### 3.2 Peptides CP and G51 form amyloid-like fibrils

The study of oligomerization of CP peptide has shown that this peptide can form aggregates after incubation for 24 hours at 55ºC in concentrations of 0.2-1 mM both in a phosphate-salt buffer (PBS, data not shown) and in 10% trifluoroethanol, which is used as membrane-microenvironment mimicking system for in-solution peptide conformation studies in some research [41–43]. Analysis of aggregates using electron and atomic force microscopy (Figure 2a and 2d) has revealed that the obtained aggregates are morphologically similar to amyloid fibrils. The use of electron microscopy provides high-resolution imaging of the object, which, in combination with atomic force microscopy that gives an idea of the average content of various structures in solution, allows for the most accurate characterization of the sample under investigation.

**Figure 1.**
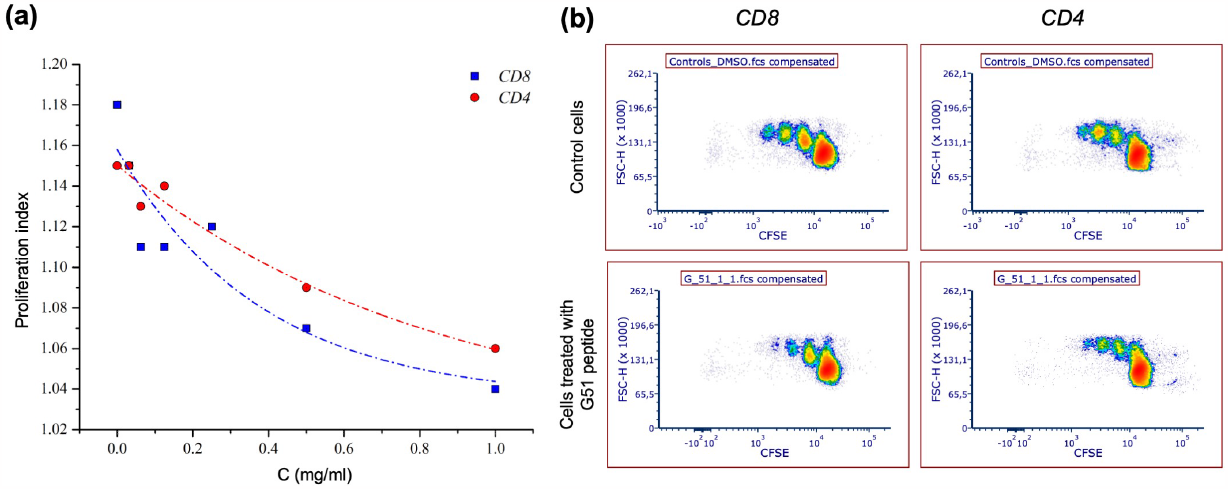
(a) Dependence of the cell proliferation index (CD8, blue and CD4, red) on the G51 peptide concentration (C, mg/ml) (b) Representative flow cytometry plots showing the proliferation of CFSE-labeled CD8 and CD4 T-cells: control ConA-stimulated cells (up) and ConA-stimulated cells in the presence of NS1-peptide.

**Figure 2.**
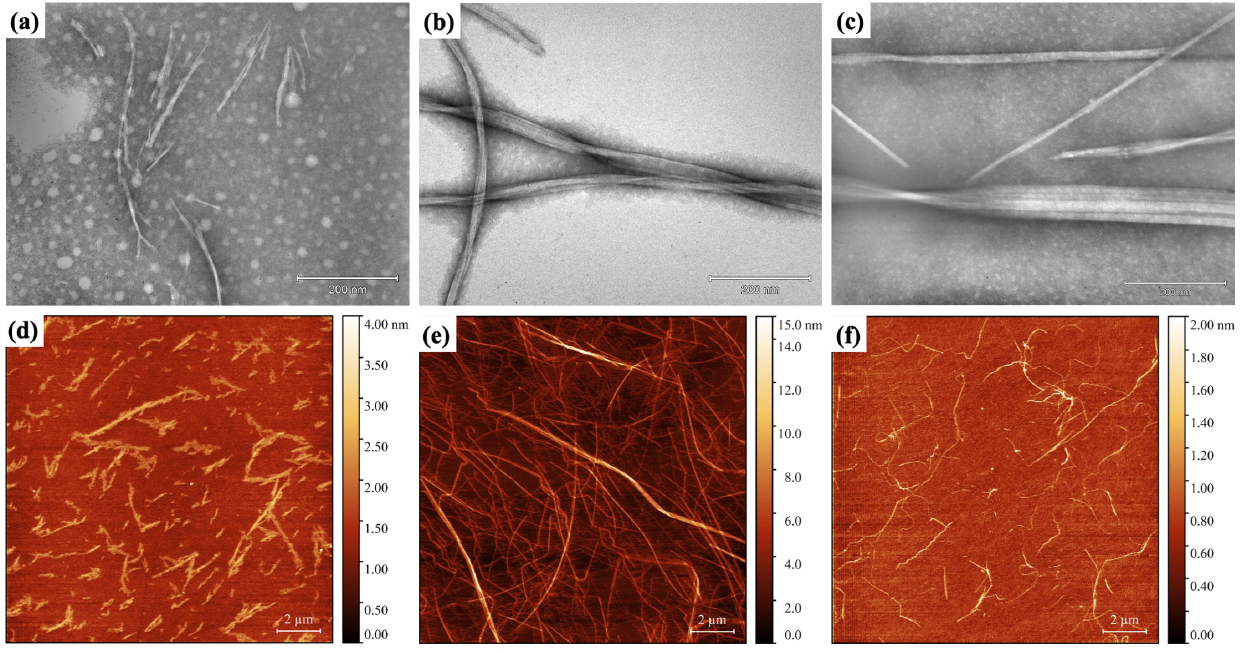
Electron microphotographs of (a) CP peptide aggregates, (b) G51 peptide aggregates, and (c) a mixture of CP and G51 peptides obtained using negative staining. The scale bar is 200 nm. Atomic force microscopy results of (d) CP peptide aggregates, (e) G51 peptide aggregates, and (f) a mixture of CP and G51 peptide aggregates. The scale bar is 2 μm, and the pseudo-color ruler on the right indicates the height of the objects (nm).

Analysis of the binding of the resulting aggregates with Congo red dye (Figure 3a) and Thioflavin T (Figure 3b,c), which are characteristic of abnormal fibrils [44,45], showed that the obtained fibrils have an amyloid-like nature. The absorption spectrum of Congo red shifted to the right, and the intensity of Thioflavin T fluorescence increased by about 30 times.

**Figure 3.**
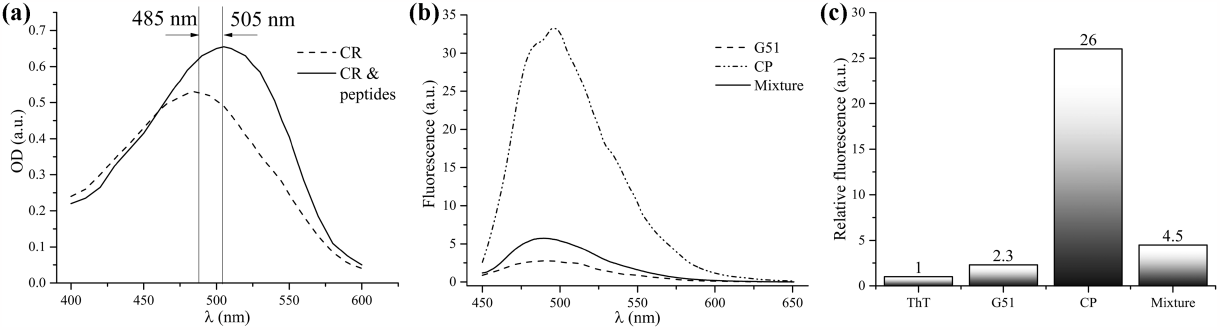
(a) Results of studying the binding of peptide fibrils with Congo red dye using optical spectroscopy. CR - Congo red dye, “CR & peptides” - characteristic curve of the dye-peptide mixture. (b) Results of studying the binding of G51 and CP peptide fibrils and their mixture with thioflavin T dye using spectrofluorimetry (“G51,” “CP,” and “Mixture” respectively). (c) Integral characteristics of the fluorescence quantum yield of G51, CP, and their mixture when bound to ThT dye (“G51,” “CP,” “Mixture” respectively; “ThT” - ThT dye solution).

These data indicate that the peptide corresponding to the core region of TCR-alpha is capable of homo-oligomerization and the formation of amyloid-like fibrils.

In experiments studying the biological activity of the NS1 protein of influenza A H1N1 strain that caused the pandemic of 1918, it was found that one of the peptides (^**140**^**rleylillraft**^**151**^, **G51**), corresponding to the primary structure of the NS1 protein fragment of influenza A virus [20] and having the highest homology with the CP peptide among the proteins of influenza A virus, is capable of oligomerization and the formation of insoluble fibrillar aggregates in concentrations of 0.2-1 mM after incubation for 24 hours at 55ºC [24].

Electron and atomic force microscopy (Figure 2b and e, respectively) have shown that the aggregates consist of fibrils. Considering that the quantum yield of Thioflavin T fluorescence (Figure 3b,c) increases by less than 2 times in the sample containing G51 peptide, it can be assumed that the number and spatial characteristics of binding sites of this dye in the fibrils of CP and G51 peptides may differ significantly.

The ability of fibrillar aggregates to alter the spectral characteristics of the specific Congo red dye (Figure 2a) allows for their amyloid-like nature.

It should be noted that according to the results of analytical gel filtration in 10% trifluoroethanol, it was determined that approximately 75% of CP and G51 peptides form aggregates with a molecular weight of more than 20 kDa after incubation for 24 hours (data not shown).

### 3.2 Mutual influence of studied peptides on the oligomerization

To investigate the mutual influence on oligomerization, a joint incubation of CP and G51 peptides was carried out in equimolar ratio (1 mM concentration per peptide) for one day at 55°C. The results of electron and atomic force microscopy (Figure 2c and 2f) showed the presence of fibrils that were significantly different in morphology from both the CP peptide fibrils and the G51 peptide fibrils.

Optical spectra of the peptide mixture with Congo red showed a shift of the absorption peak to the right, indicating the presence of amyloid-like fibrils (Figure 3a).

However, the study of Congo red binding in this modification is not a quantitative method, and the shift of the absorption peak only indicates the presence of fibrils. To compare the number of fibrils formed from the CP peptide in the presence and absence of the G51 peptide, we used the fluorescent dye thioflavin T (Figure 3b). Figure 3c shows the integral characteristics of the quantum yield of fluorescence of the CP and G51 peptides and their mixture when bound to ThT. Since the G51 peptide did not significantly increase the quantum yield of ThT upon complex formation, and there was no interference with fibrillogenesis of the peptides, the increase in fluorescence quantum yield of the fibrils in the peptide mixture should be additive, representing the sum of the quantum yields of the components of the mixture. However, during joint incubation, the quantum yield was substantially less than the sum of the increased quantum yields of the mixture components, indirectly confirming a smaller number of fibrils in the mixture observed by atomic force microscopy.

### 3.3 Fibril structural characterization

The nature of the structures formed upon dissolution of the peptides together and separately was investigated using small-angle neutron scattering. The experimental scattering curves are presented in Figure 4a (scatter plot). It was shown that when the peptides were mixed, there was a decrease in scattering intensity indicating a decrease in the concentration of scattering objects, which is consistent with the results of atomic force microscopy.

**Figure 4.**
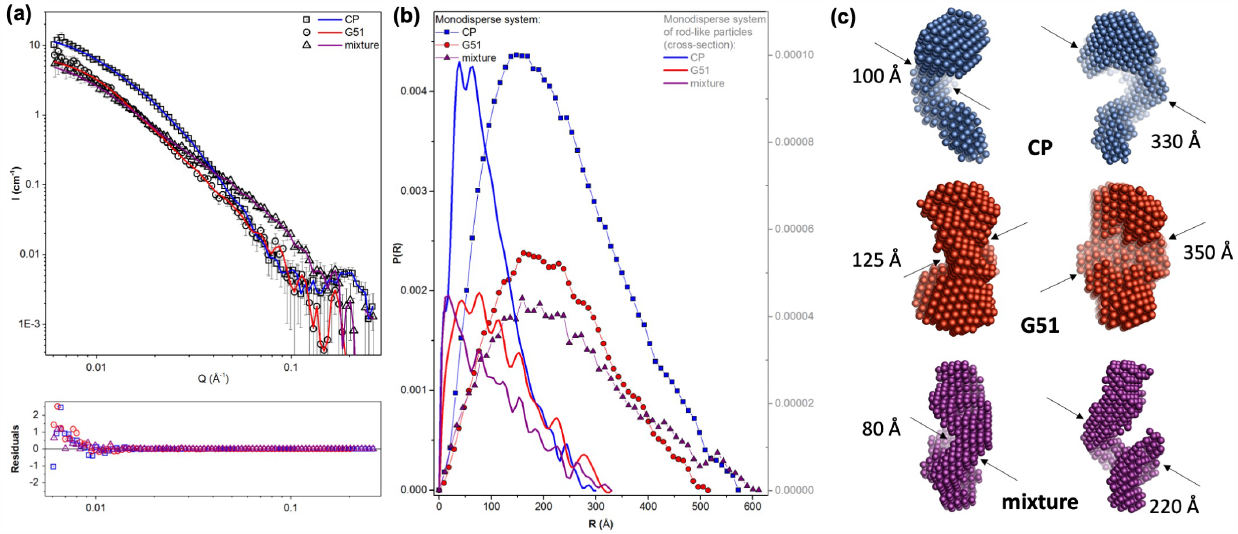
(a) SANS spectra of CP, G51 or peptide mixture samples. (b) Distance distribution function P(R) for a monodisperse system (lines) and P(R) of the cross-section assuming monodisperse system of rod-like particles (scatter and lines) of CP, G51 or peptide mixture samples, calculated based on SANS spectra. *I (cm−1*): scattering intensity, *q (Å):* magnitude of the momentum transfer, *R (Å):* distance. (c) Visualization of scattering objects, measured by SANS, using ATSAS program (CP – blue, G51 – red, mixture – purple spheres).

Curves in real space (the distance distribution function P(R) for a monodisperse system and P(R) of the cross-section assuming monodisperse system of rod-like particles) were calculated using GNOM program was used (ATSAS package) (Fig. 4b). From Figure 4b, the characteristic sizes of the scattering objects analyzed under the assumption of monodispersity of the system are similar (scatter plot), as well as R_g_ (Table 1, R_g_), while the characteristic function of pair distances for the system of rod-like particles is different for the CP peptide, G51 peptide, and their mixture (Table 1, D_max_). Using ATSAS program the probable structures of scattering objects were visualized (Fig.4c). It was shown that such objects could possess the fibril-like structure. It should be noted that the characteristic size of the peptide mixture, observed by SANS, is significantly smaller than that of the peptides separately, and the G51 peptide demonstrates a wide range of D_max_ sizes, which corresponds to the trend observed in AFM studies (Table 1, Height; Fig. 5a).

**Table 1.** Comparison of fibril parameters obtained from the SANS data (R_g_, R_c_ and D_max_) and AFM data (height), Å.

| Sample | $R_g$ ($\text{\AA}$) | $R_c$ ($\text{\AA}$) | $\sim D_{max}$ ($\text{\AA}$) | Height ($\text{\AA}$) |
| --- | --- | --- | --- | --- |
| CP | $180 \pm 12$ | $76 \pm 3$ | 50 | $11 \pm 2$ |
| G51 | $171 \pm 8$ | $94 \pm 6$ | 50-100 | $30 \pm 4$ |
| mixture | $191 \pm 7$ | $87 \pm 7$ | 20 | $7 \pm 1$ |

**Figure 5.**
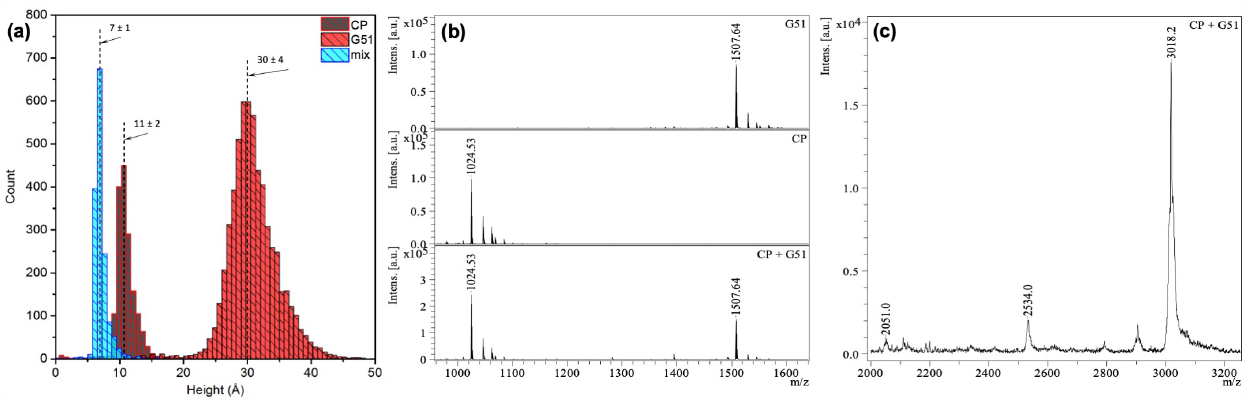
(a) Height distribution of objects calculated from atomic force microscopy results. (b) Fragment of the MALDI mass spectrum (top) of peptide CP, m/z = 1024.5; (middle) of peptide G51, m/z = 1507.6; (bottom) of their mixture in the monomer mass range. The spectrum was obtained in the reflector mode of positive ion registration. (c) Fragment of the MALDI mass spectrum of the mixture of peptides CP and G51 in the mass range of homo- and heterodimers. The spectrum was obtained in the linear mode of positive ion registration. CP homodimer – m/z = 2051, G51 homodimer – m/z = 3018, heterodimer – m/z = 2534.

Thus, based on the ThT and CR analysis, as well as SANS and AFM, it could by concluded that the G51 peptide prevented the formation of amyloid-like fibrils from the CP peptide.

Since gel-like structures were formed in both the peptide samples and the mixture after one day of incubation, such samples were unsuitable for analysis by gel filtration. Therefore, similar to [46], we used MALDI mass spectrometry to detect the presence of heterooligomers in the composition of fibrils. Figure 5b shows a fragment of the mass spectrum of the peptides separately and their mixture in the mass range corresponding to monomers. Figure 5c shows a fragment of the mass spectrum of the peptide mixture in the range corresponding to dimers.

Even though the mass spectra of fibril suspensions contain intact monomers (Figure 5b), homo- and heterodimers can be observed in the mass range corresponding to dimers (Figure 5c). The obtained results suggest that the NS1 protein fragment can prevent the oligomerization of the intramembrane domain of TCR-alpha by forming heterodimers. However, since the intensity of the peptide peaks in the spectra is not proportional to their concentration in the MALDI ionization method, it is impossible to establish the ratio of the number of homo-, heterodimers and monomers based on this mass spectrum.

### 3.4 Scheme of interaction between peptides CP and G51

We used computer modeling for the formation of homo- and hetero-oligomers of peptides G51 and CP to test the ability of peptide G51 to interact with peptide CP. For this study, our strategic approach involved initially conducting docking to select the most probable folding of two peptides, and then sequentially using docking with necessary affinity transformations to create a fibril. The obtained complexes were then subjected to molecular dynamics simulations. It turned out that both homofibrils, only from G51 or CP, and heterofibrils, from CP and G51, assembled using step-by-step docking were stable (Fig.6a) and not prone to dissociation during molecular dynamics simulations. The resulting structures were deposited [47].

**Figure 6.**
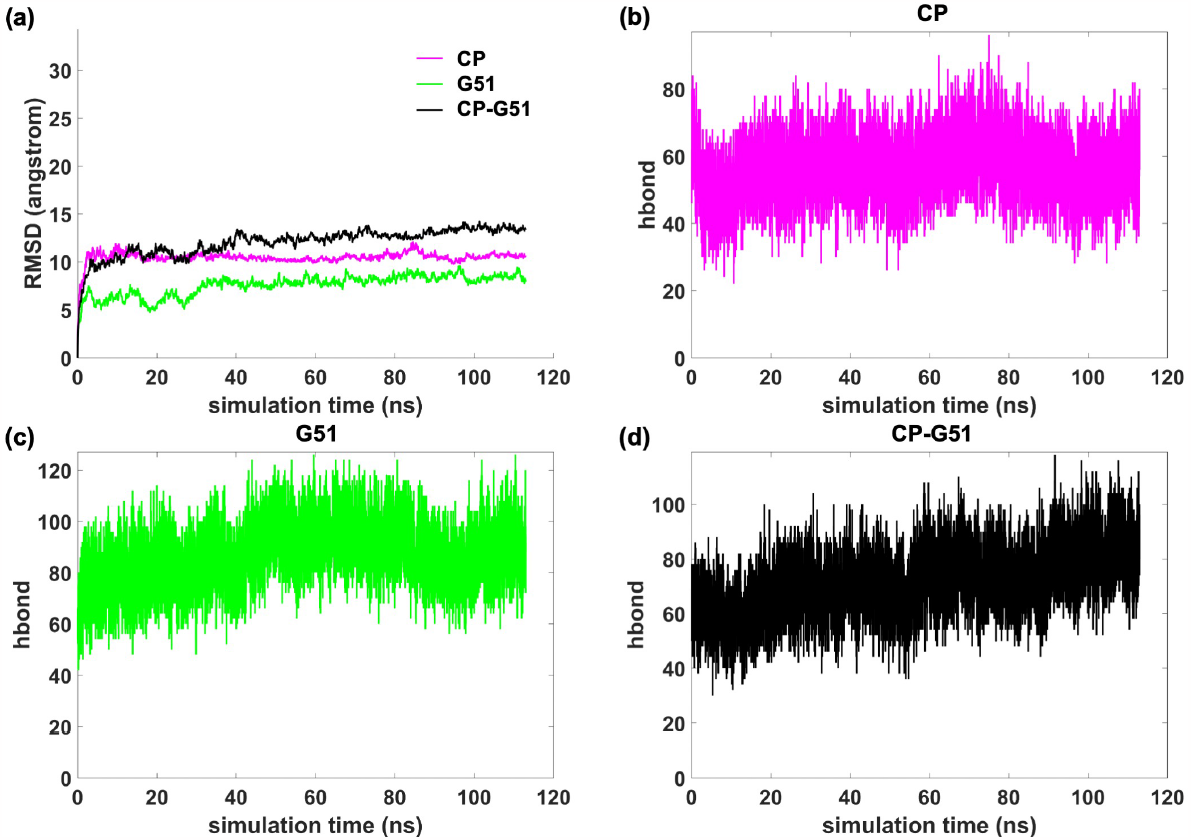
(a) RMSD. Evolution of the number of hydrogen bonds for (b) CP, (c) G51 и (d) peptide mixture.

In all three cases, stability is mainly due to powerful hydrophobic contacts and hydrogen bonds between the folded into a half ring peptide forming the fibrils (Fig. 7). In the case of CP, the monomers are positioned on top of each other so that, in addition to Leu2, Ile4, Leu5, Leu6, Leu7, and Val9 which are localized on top of each other and form hydrophobic contacts, Leu2, Leu5, and Val9 also interact with each other. The positive charge of the side amino group Lys8 is compensated by an ionic and strong hydrogen bond with the negatively charged terminal carboxyl group Val9.

**Figure 7.**
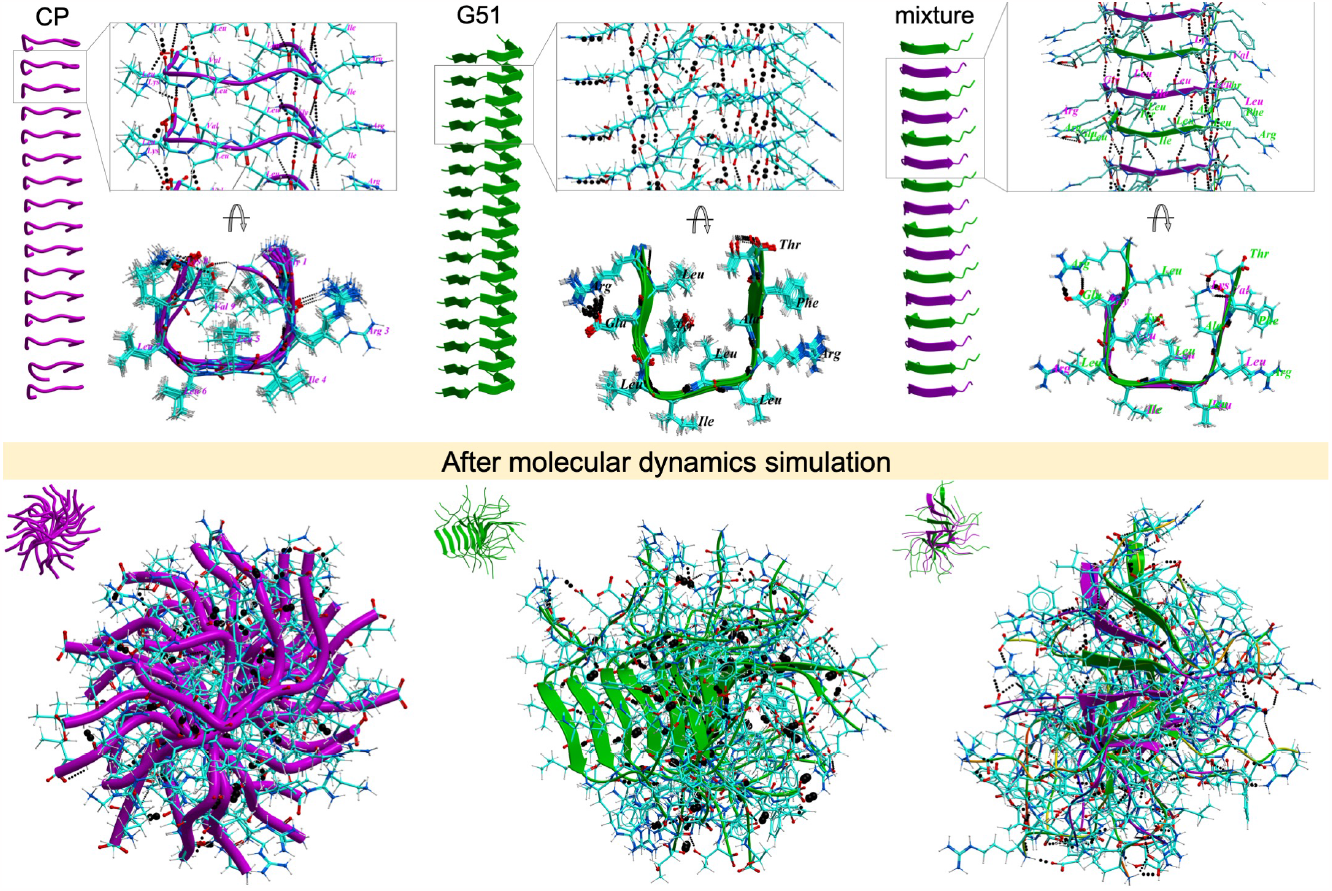
Rendering only (side view), a piece of fibril with hydrogen bonds, structure with rendering and amino acid names (top view). Atoms are represented by the following colours: carbon – cyan, oxygen – red, nitrogen – blue. Hydrogen bonds are shown as dashed lines; the thickness depends on the binding energy.

In the case of G51 (Fig.7, central column, inset), the packing in the fibril is organized in such a way that the main Van-der-Waals contribution to free energy is provided by Leu2, Tyr4, Leu5, Ile6, Leu7, Leu8, Val9, and Phe11 which are located on top of each other. Additional contributions to the hydrophobic component of free energy in G51 are also made by Van der Waals interactions between the side groups of Leu2, Tyr4, and the methyl group of Thr12; the side groups of Leu2, Tyr4, and Ala10; the side group of Ala10 and the methyl group of Thr12. The positively charged guanidine group Arg1 forms an ionic and strong hydrogen bond with the negatively charged side carboxyl group Glu3. As in CP, the folding of monomers on top of each other in G51 occurs in such a way that a network of hydrogen bonds is formed between the peptide backbones as in beta-sheets: the peptide carbonyl oxygen acts as an acceptor for the neighboring peptide amino group.

In the case of the CP-G51 mixture (Fig.7, right column, inset), depending on the conditions, mixed fibrils can be formed by overlapping CP and G51 on top of each other, as well as complexes of docked fibrils consisting only of CP or only of G51. In the case of mixed fibrils, both CP and G51 are folded into semi-circles, with the side groups of CP peptides adjacent to the side groups of G51 peptides as follows: CP/Leu2 – G51/Tyr4, CP/Arg3 – G51/Leu5, CP/Ile4 – G51/Ile6, CP/Leu5 – G51/Leu7, CP/Leu6 – G51/Leu8, CP/Leu7 – G51/Arg9, CP/Lys8 – G51/Ala10, and CP/Val9 – G51/Phe11. Due to the semi-circular folding, additional contributions to the hydrophobic component of free energy are made by Van-der-Waals interactions located inside the semicircles G51/Leu2, G51/Tyr4, G51/Leu7, G51/Leu2, G51/Ala10, CP/Leu2, and CP/Leu5. As in the case of fibrils consisting only of CP or only of G51, peptide backbones in mixed fibrils also form a network of hydrogen bonds as in beta-sheets. In addition to these hydrogen bonds, hydrogen bonds are also formed between the charged terminal groups and NH and O located near them in peptide bonds. As in homogeneous fibrils of CP and G51, in mixed fibrils, the positive charge of the side amino group Lys8 in CP is compensated by an ionic and strong hydrogen bond with the negatively charged terminal carboxyl group Val9, while in G51, the positively charged guanidine group Arg1 forms an ionic and strong hydrogen bond with the negatively charged side carboxyl group Glu3.

Analysis of the dynamics of the number of hydrogen bonds (Fig.6b-d), as well as the data presented in Table S1, which shows the percentage of snapshots with specific hydrogen bonds, showed that in all studied cases, the most stable hydrogen bonds were those connecting the central peptide fragments. This fact is consistent with experimental data indicating a wide range of fibril widths.

Analysis of the evolution of the values of contributions to the free energy presented in Fig. S1. showed that the formation of fibrils from the individual peptides under study is contributed by the Coulombic component, which is caused by both strong hydrogen-ionic bonds at the ends and hydrogen bonds between the peptide backbones.

During the dynamics, in all the cases considered, twisting of the fibrils occurred, which is also consistent with electron microscopy data.

In cases where solutions of pre-formed homogeneous CP and G51 fibrils are mixed, fibril junctions may occur. The mutual arrangement of CP and G51 fibrils relative to each other is such that fragments carrying opposite charges localize next to each other.

Based on the obtained data, it can be suggested that peptide G51 stabilizes CP oligomers in a state different from when the latter is present in homo-oligomers.

## 4. Discussion

Peptides CP and G51 individually tend to form beta structures and amyloid-like fibrils. However, in a mixture, in the presence of a partner, peptides apparently form heterodimers by interacting with opposite charged amino acids and hydrophobic regions, resulting in a decrease in the quantum yield of thioflavin T fluorescence and also a reduction in the number of fibrils.

The ability to form amyloid-like fibrils by the TCR fragment in a model membrane environment may be related to the peculiarities of signal transduction in cells. Molecular mechanisms that allow regulation of signal transduction by proteins at low concentrations are well known. One such mechanism is the prion-like mechanism of inducing conformational transitions [48]. In this mechanism of regulation, a molecule in a special prion-like conformation induces a change in the spatial structure of the cellular factor, thus involving it in further conformational transitions in a chain reaction. Oligomerization-induced activation of T-cell receptors of various types has been demonstrated quite some time ago [49], but data on the formation of amyloid-like oligomers *in vivo* is absent in the literature.

The discovered ability of the NS1 protein fragment to fibrillogenicity may lead to the possibility of recruiting cellular factors involved in antiviral response and thus play an important role in increasing the pathogenicity of influenza virus. For example, it is known that in the process of interaction of respiratory syncytial virus with a cell, the NS1 protein undergoes structural changes and acquires amyloid-like properties, which determine its ability to interact with many cellular factors [50,51].

Although the mechanism of enhancing the response to viral infection through conformational transmission is involved in RIG-1/MAVS-mediated antiviral response [52], the involvement of NS1 in activating or inhibiting this pathway has not been demonstrated.

The NS1 protein of the influenza virus is prone to structural polymorphism, which also provides its multifunctionality [51,53], but its ability to form amyloid-amyloidlike fibrils *in vivo* has not yet been demonstrated. Previous data suggest that the 140-151 aa region of NS1 may serve as a potential amyloidogenic determinant of this protein [24].

Usually, the induction of conformational transition requires homology between the protein sequences in the prion-like conformation and the effector protein [54]. In this case, if the viral protein can acquire a prion-like conformation and possesses an amyloidogenic determinant homologous to a cellular factor, induction of the conformational transition of the cellular factor can occur, recruiting it into prion-like oligomers. On the other hand, in some cases, polypeptides that are close in primary structure can prevent each other’s oligomerization and fibrillogenesis. For example, in [55] it was shown that a peptide containing the fibrillogenic determinant of insulin and additional arginine residues prevents fibrillogenesis of this protein. In another study, it was shown that transthyretin with altered amino acid sequence can prevent fibrillogenesis of the same protein with another amino acid substitution [56]. Furthermore, there are instances where amyloid-like aggregates of viral proteins are able to affect the physiological and pathological aggregation of human proteins [57].

Therefore, in the presence of interaction between CP and G51 peptides, both scenarios of fibrillogenesis induction and oligomerization inhibition could be realized. The inhibition of oligomerization of the intramembrane domain of TCR by NS1 protein, that we discovered in the peptide model *in vitro*, may lead to the blocking of T-cell immune response and immunosuppression during influenza. This interaction could serve as a promising target for drug discovery. Furthermore, due to the strong evolutionary pressure leading to the emergence of resistance of current viruses to direct antiviral drugs and vaccines [58], the most promising avenues involve the development of drugs aimed either at the metabolic pathways of the host or at the interface between virus and host proteins.

## Conclusion

It has been shown that a peptide corresponding to the primary structure of the potential immunosuppressive domain (ISD) of the NS1 protein of the influenza virus is capable of inhibiting PBMC cell proliferation induced by concanavalin A.

The use of electron atomic force microscopy, spectrophotometry with Congo red dye, and fluorimetry with thioflavin T dye demonstrated that the T-cell receptor core peptide (CP) and its homologous peptide G51 from the NS1 influenza virus (as was shown by us earlier [24]) can form amyloid-like fibrils *in vitro*. Additionally, peptide G51 can form heterooligomers with peptide CP and modulate its oligomerization. Based on hypothesis that in nature, two peptides join together, and the resulting structure acts as a seed for the subsequent growth of the fibril, we aimed to mimic the natural assembly process using computer modeling. Our approach enabled us to obtain probable fibril conformations and describe the molecular interactions that stabilize the formed fibril. Results from computer modeling suggest that the homology of the primary structures of model peptides may play a role in this interaction, and indicate potential molecular mechanisms for their specific interaction. The obtained data from the *in vitro* model peptide system suggest the presence of a functional interaction between the NS1 fragment of the influenza A virus and the intramembrane domain of the alpha-subunit of the T-cell receptor.

The hypothesized scenario based on the obtained data is as follows: a fragment of the NS1 protein, homologous to the G51 peptide, when presented to the MHC for T cells, is capable of acting on the transmembrane domain of the T cell receptor through a prion-like mechanism, inactivating normal signal transmission and stimulation of T cell division.

Molecular modeling has suggested molecular structures of NS1 and CP that are critical for this interaction. Combined with the capabilities of modern molecular modeling (similar to [59]), our proposed description of the potential structural mechanism of immunosuppression provides the basis for the design of small-molecule blocking drugs that allow testing the hypothesis in a cellular system. In addition, the *in vitro* model and cellular systems described in this work are suitable for searching for such potential compounds. The compounds found in this way can act as potential pharmacological substances affecting immunosuppression in influenza.

Despite extensive evidence regarding the role of influenza NS1 protein in immunosuppression, we are the first to demonstrate action in cells and propose a potential mechanism for immunosuppression of a peptide homologous to the NS1 fragment. The participation in this process of amyloid-like aggregates, capable of acting in low concentrations due to cooperativity and prion-like transfer of conformation, suggests the possibility of another fundamental process associated with functional amyloids.

## Supporting information

Supplementary Fugue S1

## Acknowledgments

The authors thanks Prof., Dr., Acad. Oleg Kisilev for generation of concept and fruitful discussion of the results.

## References

[1] R.M. Krug, Functions of the influenza A virus NS1 protein in antiviral defense, Curr Opin Virol. 12 (2015) 1–6. 10.1016/j.coviro.2015.01.007.

[2] S.N. Thulasi Raman, Y. Zhou, Networks of Host Factors that Interact with NS1 Protein of Influenza A Virus, Front Microbiol. 7 (2016) 654. 10.3389/fmicb.2016.00654.

[3] K. Das, J.M. Aramini, L.-C. Ma, R.M. Krug, E. Arnold, Structures of influenza A proteins and insights into antiviral drug targets, Nat Struct Mol Biol. 17 (2010) 530–8. 10.1038/nsmb.1779.

[4] D. Marc, Influenza virus non-structural protein NS1: interferon antagonism and beyond, Journal of General Virology. 95 (2014) 2594–2611. 10.1099/vir.0.069542-0.

[5] R. Rajsbaum, R.A. Albrecht, M.K. Wang, N.P. Maharaj, G.A. Versteeg, E. Nistal-Villán, A. García-Sastre, M.U. Gack, Species-Specific Inhibition of RIG-I Ubiquitination and IFN Induction by the Influenza A Virus NS1 Protein, PLoS Pathog. 8 (2012) e1003059. 10.1371/journal.ppat.1003059.

[6] J. Rehwinkel, C.P. Tan, D. Goubau, O. Schulz, A. Pichlmair, K. Bier, N. Robb, F. Vreede, W. Barclay, E. Fodor, C. Reis e Sousa, RIG-I Detects Viral Genomic RNA during Negative-Strand RNA Virus Infection, Cell. 140 (2010) 397–408. 10.1016/j.cell.2010.01.020.

[7] J.-Y. Min, S. Li, G.C. Sen, R.M. Krug, A site on the influenza A virus NS1 protein mediates both inhibition of PKR activation and temporal regulation of viral RNA synthesis, Virology. 363 (2007) 236–43. 10.1016/j.virol.2007.01.038.

[8] N.C. Robb, E. Fodor, The accumulation of influenza A virus segment 7 spliced mRNAs is regulated by the NS1 protein, J Gen Virol. 93 (2012) 113–8. 10.1099/vir.0.035485-0.

[9] P. Fortes, A. Beloso, J. Ortín, Influenza virus NS1 protein inhibits pre-mRNA splicing and blocks mRNA nucleocytoplasmic transport, EMBO J. 13 (1994) 704–12. http://www.ncbi.nlm.nih.gov/pubmed/8313914 x(accessed July 24, 2017).

[10] A. Mor, A. White, K. Zhang, M. Thompson, M. Esparza, R. Muñoz-Moreno, K. Koide, K.W. Lynch, A. García-Sastre, B.M.A. Fontoura, Influenza virus mRNA trafficking through host nuclear speckles, Nat Microbiol. 1 (2016) 16069. 10.1038/nmicrobiol.2016.69.

[11] W. Nacken, A. Schreiber, D. Masemann, S. Ludwig, The Effector Domain of the Influenza A Virus Nonstructural Protein NS1 Triggers Host Shutoff by Mediating Inhibition and Global Deregulation of Host Transcription When Associated with Specific Structures in the Nucleus, MBio. 12 (2021). 10.1128/mBio.02196-21.

[12] I. Marazzi, J.S.Y. Ho, J. Kim, B. Manicassamy, S. Dewell, R.A. Albrecht, C.W. Seibert, U. Schaefer, K.L. Jeffrey, R.K. Prinjha, K. Lee, A. García-Sastre, R.G. Roeder, A. Tarakhovsky, Suppression of the antiviral response by an influenza histone mimic, Nature 2012 483:7390. 483 (2012) 428–433. 10.1038/nature10892.

[13] S. Qin, Y. Liu, W. Tempel, M.S. Eram, C. Bian, K. Liu, G. Senisterra, L. Crombet, M. Vedadi, J. Min, Structural basis for histone mimicry and hijacking of host proteins by influenza virus protein NS1, Nat Commun. 5 (2014) 3952. 10.1038/ncomms4952.

[14] L. Zhu, J. Qin, A Viral Protein Mimics Histone to Hijack Host MORC3, Structure. 27 (2019) 883–885. 10.1016/j.str.2019.05.007.

[15] Y. Genzel, C. Dietzsch, E. Rapp, J. Schwarzer, U. Reichl, MDCK and Vero cells for influenza virus vaccine production: a one-to-one comparison up to lab-scale bioreactor cultivation, Appl Microbiol Biotechnol. 88 (2010) 461–475. 10.1007/s00253-010-2742-9.

[16] S.K. Dankar, E. Miranda, N.E. Forbes, M. Pelchat, A. Tavassoli, M. Selman, J. Ping, J. Jia, E.G. Brown, Influenza A/Hong Kong/156/1997(H5N1) virus NS1 gene mutations F103L and M106I both increase IFN antagonism, virulence and cytoplasmic localization but differ in binding to RIG-I and CPSF30, Virol J. 10 (2013) 243. 10.1186/1743-422X-10-243.

[17] R. Volmer, B. Mazel-Sanchez, C. Volmer, S.M. Soubies, J.-L. Guérin, Nucleolar localization of influenza A NS1: striking differences between mammalian and avian cells, Virol J. 7 (2010) 63. 10.1186/1743-422X-7-63.

[18] B. Nicholas, A. Bailey, K.J. Staples, T. Wilkinson, T. Elliott, P. Skipp, Immunopeptidomic analysis of influenza A virus infected human tissues identifies internal proteins as a rich source of HLA ligands, PLoS Pathog. 18 (2022) e1009894. 10.1371/JOURNAL.PPAT.1009894.

[19] M.E. Call, K.W. Wucherpfennig, J.J. Chou, The structural basis for intramembrane assembly of an activating immunoreceptor complex, Nat Immunol. 11 (2010) 1023–9. 10.1038/ni.1943.

[20] O.I. Kiselev, Human endogenous retroviruses, placenta and immunosuppression, “Rostok,” St. Petersburg, 2014.

[21] M. Ali, M.R.R. De Planque, N.T. Huynh, N. Manolios, F. Separovic, Biophysical studies of a transmembrane peptide derived from the T cell antigen receptor, Letters in Peptide Science. 8 (2001) 227–233. 10.1007/BF02446521.

[22] G. Zheng, A.M. Torres, M. Ali, N. Manolios, W.S. Price, NMR study of the structure and self-association of core peptide in aqueous solution and DPC micelles, Biopolymers. 96 (2011) 177–180. 10.1002/bip.21423.

[23] V. Bender, M. Ali, M. Amon, E. Diefenbach, N. Manolios, T Cell Antigen Receptor Peptide-Lipid Membrane Interactions Using Surface Plasmon Resonance, Journal of Biological Chemistry. 279 (2004) 54002–54007. 10.1074/jbc.M403909200.

[24] A.A. Shaldzhyan, Y.A. Zabrodskaya, I.L. Baranovskaya, M.V. Sergeeva, A.N. Gorshkov, I.I. Savin, S.M. Shishlyannikov, E.S. Ramsay, A.V. Protasov, A.P. Kukhareva, V.V. Egorov, Old dog, new tricks: Influenza A virus NS1 and in vitro fibrillogenesis, Biochimie. 190 (2021) 50–56. 10.1016/j.biochi.2021.07.005.

[25] Y. Zabrodskaya, M. Plotnikova, N. Gavrilova, A. Lozhkov, S. Klotchenko, A. Kiselev, V. Burdakov, E. Ramsay, L. Purvinsh, M. Egorova, V. Vysochinskaya, I. Baranovskaya, A. Brodskaya, R. Povalikhin, A. Vasin, Exosomes Released by Influenza-Virus-Infected Cells Carry Factors Capable of Suppressing Immune Defense Genes in Naïve Cells, Viruses 2022, Vol. 14, Page 2690. 14 (2022) 2690. 10.3390/V14122690.

[26] S. Rastogi, V. Sharma, P.S. Bharti, K. Rani, G.P. Modi, F. Nikolajeff, S. Kumar, The Evolving Landscape of Exosomes in Neurodegenerative Diseases: Exosomes Characteristics and a Promising Role in Early Diagnosis, International Journal of Molecular Sciences 2021, Vol. 22, Page 440. 22 (2021) 440. 10.3390/IJMS22010440.

[27] D. Nečas, P. Klapetek, Gwyddion: an open-source software for SPM data analysis, Open Physics. 10 (2012) 181–188. 10.2478/s11534-011-0096-2.

[28] A.I. Kuklin, A.Kh. Islamov, V.I. Gordeliy, Scientific Reviews: Two-Detector System for Small-Angle Neutron Scattering Instrument, Neutron News. 16 (2005) 16–18. 10.1080/10448630500454361.

[29] A.I. Kuklin, D. V Soloviov, A. V Rogachev, P.K. Utrobin, Y.S. Kovalev, M. Balasoiu, O.I. Ivankov, A.P. Sirotin, T.N. Murugova, T.B. Petukhova, Y.E. Gorshkova, R. V Erhan, S.A. Kutuzov, A.G. Soloviev, V.I. Gordeliy, New opportunities provided by modernized small-angle neutron scattering two-detector system instrument (YuMO), J Phys Conf Ser. 291 (2011) 012013. 10.1088/1742-6596/291/1/012013.

[30] A.G. Soloviev, T.M. Solovjeva, O.I. Ivankov, D. V Soloviov, A. V Rogachev, A.I. Kuklin, SAS program for two-detector system: seamless curve from both detectors, J Phys Conf Ser. 848 (2017) 012020. 10.1088/1742-6596/848/1/012020.

[31] D.I. Svergun, Determination of the regularization parameter in indirect-transform methods using perceptual criteria, J Appl Crystallogr. 25 (1992) 495–503. 10.1107/S0021889892001663.

[32] D.I. Svergun, Restoring Low Resolution Structure of Biological Macromolecules from Solution Scattering Using Simulated Annealing, Biophys J. 76 (1999) 2879–2886. 10.1016/S0006-3495(99)77443-6.

[33] Y.A. Zabrodskaya, V. V. Egorov, A. V. Sokolov, A. V. Shvetsov, Y.E. Gorshkova, O.I. Ivankov, V.A. Kostevich, N.P. Gorbunov, E.S. Ramsay, N.D. Fedorova, A.B. Bondarenko, V.B. Vasilyev, Caught red handed: modeling and confirmation of the myeloperoxidase ceruloplasmin alpha-thrombin complex, BioMetals. 35 (2022) 1157–1168. 10.1007/s10534-022-00432-2.

[34] R. Abagyan, M. Totrov, D. Kuznetsov, ICM-A new method for protein modeling and design: Applications to docking and structure prediction from the distorted native conformation, J Comput Chem. 15 (1994) 488–506. 10.1002/jcc.540150503.

[35] Y.A. Arnautova, A. Jagielska, H.A. Scheraga, A New Force Field (ECEPP-05) for Peptides, Proteins, and Organic Molecules, J Phys Chem B. 110 (2006) 5025–5044. 10.1021/jp054994x.

[36] D.A. Case, K. Belfon, I.Y. Ben-Shalom, S.R. Brozell, D.S. Cerutti, T.E.I. Cheatham, V.W.D. Cruzeiro, T.A. Darden, R.E. Duke, G. Giambasu, M.K. Gilson, H. Gohlke, A.W. Goetz, R. Harris, S. Izadi, K. Kasavajhala, A. Kovalenko, R. Krasny, T. Kurtzman, T.S. Lee, S. LeGrand, P. Li., C. Lin, J. Liu, T. Luchko, R. Luo, V. Man, K.M. Merz, Y. Miao, O. Mikhailovskii, G. Monard, H. Nguyen, A. Onufriev, F. Pan, S. Pantano, R. Qi, D.R. Roe, A. Roitberg, C. Sagui, S. Schott-Verdugo, J. Shen, C.L. Simmerling, N. Skrynnikov, J. Smith, J. Swails, R.C. Walker, J. Wang, L. Wilson, R.M. Wolf, X. Wu, D.M. York, P.A. Kollman, AMBER 2020, (2020). https://ambermd.org/index.php (accessed March 6, 2023).

[37] J.A. Maier, C. Martinez, K. Kasavajhala, L. Wickstrom, K.E. Hauser, C. Simmerling, ff14SB: Improving the Accuracy of Protein Side Chain and Backbone Parameters from ff99SB, J Chem Theory Comput. 11 (2015) 3696–3713. 10.1021/ACS.JCTC.5B00255/SUPPL_FILE/CT5B00255_SI_001.PDF.

[38] A. Onufriev, D.A. Case, D. Bashford, Effective Born radii in the generalized Born approximation: The importance of being perfect, J Comput Chem. 23 (2002) 1297–1304. 10.1002/JCC.10126.

[39] B. Carrillo, J.-M. Choi, Z.A. Bornholdt, B. Sankaran, A.P. Rice, B.V. V. Prasad, The Influenza A Virus Protein NS1 Displays Structural Polymorphism, J Virol. 88 (2014) 4113–4122. 10.1128/jvi.03692-13.

[40] Y. Chen, Y. Zhu, X. Li, W. Gao, Z. Zhen, D. Dong, B. Huang, Z. Ma, A. Zhang, X. Song, Y. Ma, C. Guo, F. Zhang, Z. Huang, Cholesterol inhibits TCR signaling by directly restricting TCR-CD3 core tunnel motility, Mol Cell. 82 (2022) 1278–1287.e5. 10.1016/j.molcel.2022.02.017.

[41] H. Sticht, P. Bayer, D. Willbold, S. Dames, C. Hilbich, K. Beyreuther, R.W. Frank, P. Rösch, Structure of amyloid A4-(1-40)-peptide of Alzheimer’s disease, Eur J Biochem. 233 (1995) 293–8.

[42] W. Liu, J.M. Prausnitz, H.W. Blanch, Amyloid Fibril Formation by Peptide LYS (11-36) in Aqueous Trifluoroethanol, Biomacromolecules. 5 (2004) 1818–1823. 10.1021/bm049841e.

[43] K.H. Ruan, P. Li, R.J. Kulmacz, K.K. Wu, Characterization of the structure and membrane interaction of NH2-terminal domain of thromboxane A2 synthase., Journal of Biological Chemistry. 269 (1994) 20938–20942. 10.1016/S0021-9258(17)31912-9.

[44] H. LeVine, Thioflavine T interaction with synthetic Alzheimer’s disease beta-amyloid peptides: detection of amyloid aggregation in solution, Protein Sci. 2 (1993) 404–10. 10.1002/pro.5560020312.

[45] O.I. Antimonova, N.A. Grudinina, V. V. Egorov, D.S. Polyakov, V. V. Il’in, M.M. Shavlovskii, Interaction of the dye Congo red with fibrils of lysozyme, beta2-microglobulin, and transthyretin, Cell Tissue Biol. 10 (2016) 468–475. 10.1134/S1990519X1606002X.

[46] V. V Egorov, Y.A. Zabrodskaya, D. V Lebedev, A.N. Gorshkov, A.I. Kuklin, Structural features of the ionic self-complementary amyloidogenic peptide, J Phys Conf Ser. 848 (2017) 012022. 10.1088/1742-6596/848/1/012022.

[47] V. Tsvetkov, V. Egorov, Y. Zabrodskaya, Results of molecular dynamics of fibrils, formed by CP and G51 peptides, Zenodo. (2023). 10.5281/ZENODO.10410486.

[48] P. Tompa, The principle of conformational signaling, Chem Soc Rev. 45 (2016) 4252–84. 10.1039/c6cs00011h.

[49] M.M. Davis, Z. Reich, J.J. Boniface, D.S. Lyons, N. Borochov, E.J. Wachtel, Ligand-specific oligomerization of T-cell receptor molecules, Nature. 387 (1997) 617–620. 10.1038/42500.

[50] E. Pretel, I.E. Sánchez, M. Fassolari, L.B. Chemes, G. de Prat-Gay, Conformational Heterogeneity Determined by Folding and Oligomer Assembly Routes of the Interferon Response Inhibitor NS1 Protein, Unique to Human Respiratory Syncytial Virus, Biochemistry. 54 (2015) 5136–5146. 10.1021/acs.biochem.5b00615.

[51] E. Pretel, G. Camporeale, G. de Prat-Gay, The Non-Structural NS1 Protein Unique to Respiratory Syncytial Virus: A Two-State Folding Monomer in Quasi-Equilibrium with a Stable Spherical Oligomer, PLoS One. 8 (2013) e74338. 10.1371/journal.pone.0074338.

[52] B. Wu, S. Hur, How RIG-I like receptors activate MAVS, Curr Opin Virol. 12 (2015) 91–98. 10.1016/j.coviro.2015.04.004.

[53] B. Carrillo, J.-M. Choi, Z.A. Bornholdt, B. Sankaran, A.P. Rice, B.V.V. Prasad, The influenza A virus protein NS1 displays structural polymorphism, J Virol. 88 (2014) 4113–22. 10.1128/JVI.03692-13.

[54] M.R.H. Krebs, L.A. Morozova-Roche, K. Daniel, C. V. Robinson, C.M. Dobson, Observation of sequence specificity in the seeding of protein amyloid fibrils, Protein Sci. 13 (2004) 1933–1938. 10.1110/ps.04707004.

[55] T.J. Gibson, R.M. Murphy, Inhibition of insulin fibrillogenesis with targeted peptides, Protein Sci. 15 (2006) 1133–41. 10.1110/ps.051879606.

[56] P. Hammarström, F. Schneider, J.W. Kelly, Trans-suppression of misfolding in an amyloid disease, Science. 293 (2001) 2459–62. 10.1126/science.1062245.

[57] A. Ghosh, A.S. Pithadia, J. Bhat, S. Bera, A. Midya, C.A. Fierke, A. Ramamoorthy, A. Bhunia, Self-Assembly of a Nine-Residue Amyloid-Forming Peptide Fragment of SARS Corona Virus E-Protein: Mechanism of Self Aggregation and Amyloid-Inhibition of hIAPP, Biochemistry. 54 (2015) 2249–2261. 10.1021/acs.biochem.5b00061.

[58] A.M. Rabie, The Informative Nature of the Disappeared SARS-CoV-2 Genomic Sequences: A Mini-review with Perspectives, Advanced Chemicobiology Research. 1 (2022) 58–64. 10.37256/acbr.1220221403.

[59] A.M. Rabie, Teriflunomide: A possible effective drug for the comprehensive treatment of COVID-19, Current Research in Pharmacology and Drug Discovery. 2 (2021) 100055. 10.1016/j.crphar.2021.100055.

