## Supplementary Fugue S1 for "How the immune mousetrap works: structural evidence for the immunomodulatory action of a peptide from influenza NS1 protein"

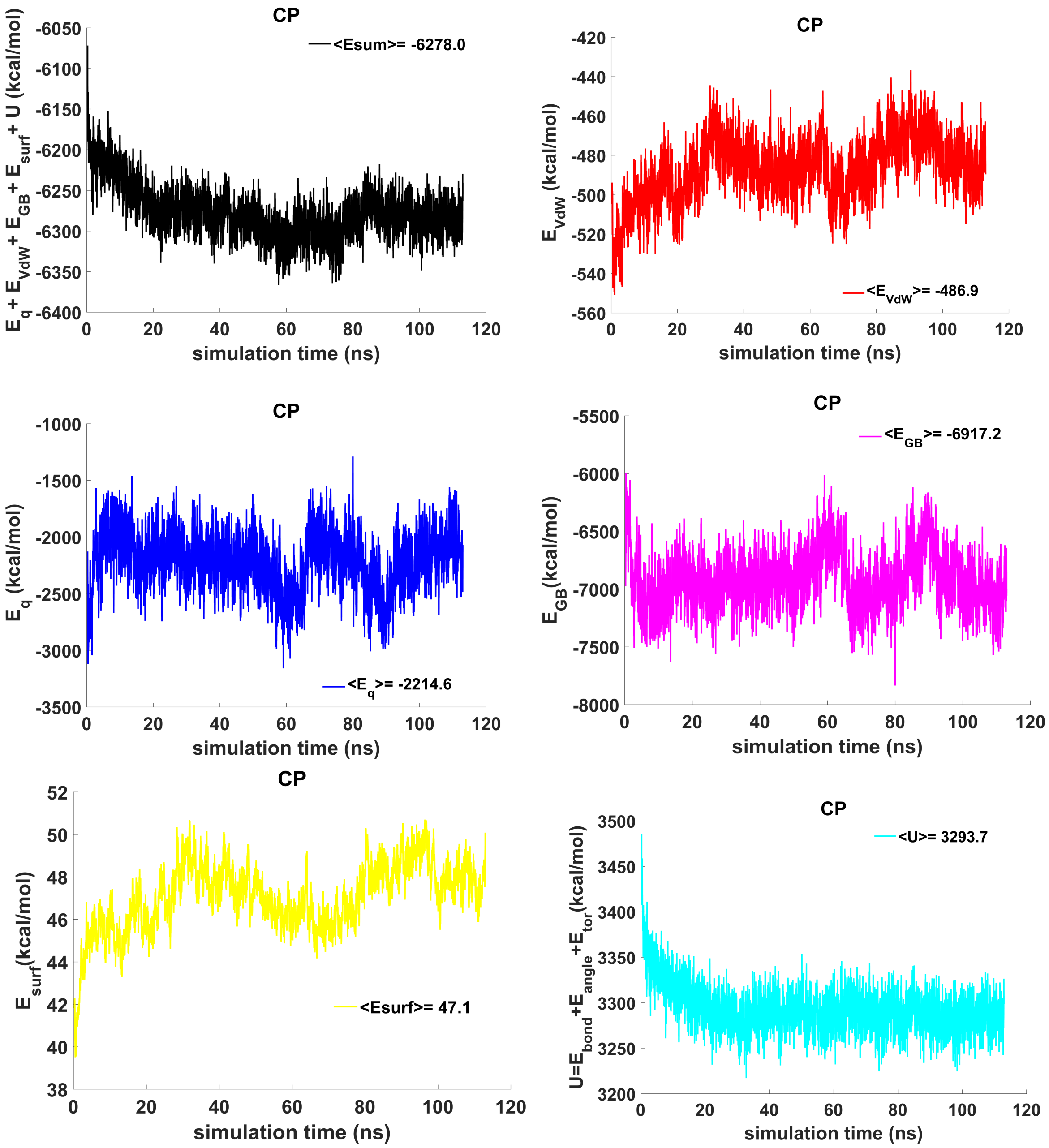


Fig.S1. A. **The contributions to free energy during MD calculations for the CP model**. $E_{eq}$ – electrostatic, $E_{vdw}$ - Van der Waals, $E_{GB}$ - polar energy of solvation, $E_{surf}$ - non-polar energy of solvation due to the hydrophobic surface available to the solvent, U = $E_{bond}$+ $E_{angle}$+ $E_{tor}$ , e.g. $E_{bond}$, $E_{angle}$ and $E_{tor}$ –bond, angle and torsion stress energies. The energy plots were smoothed using moving average method (span = 5). Average energy values are indicated in the figure legends.

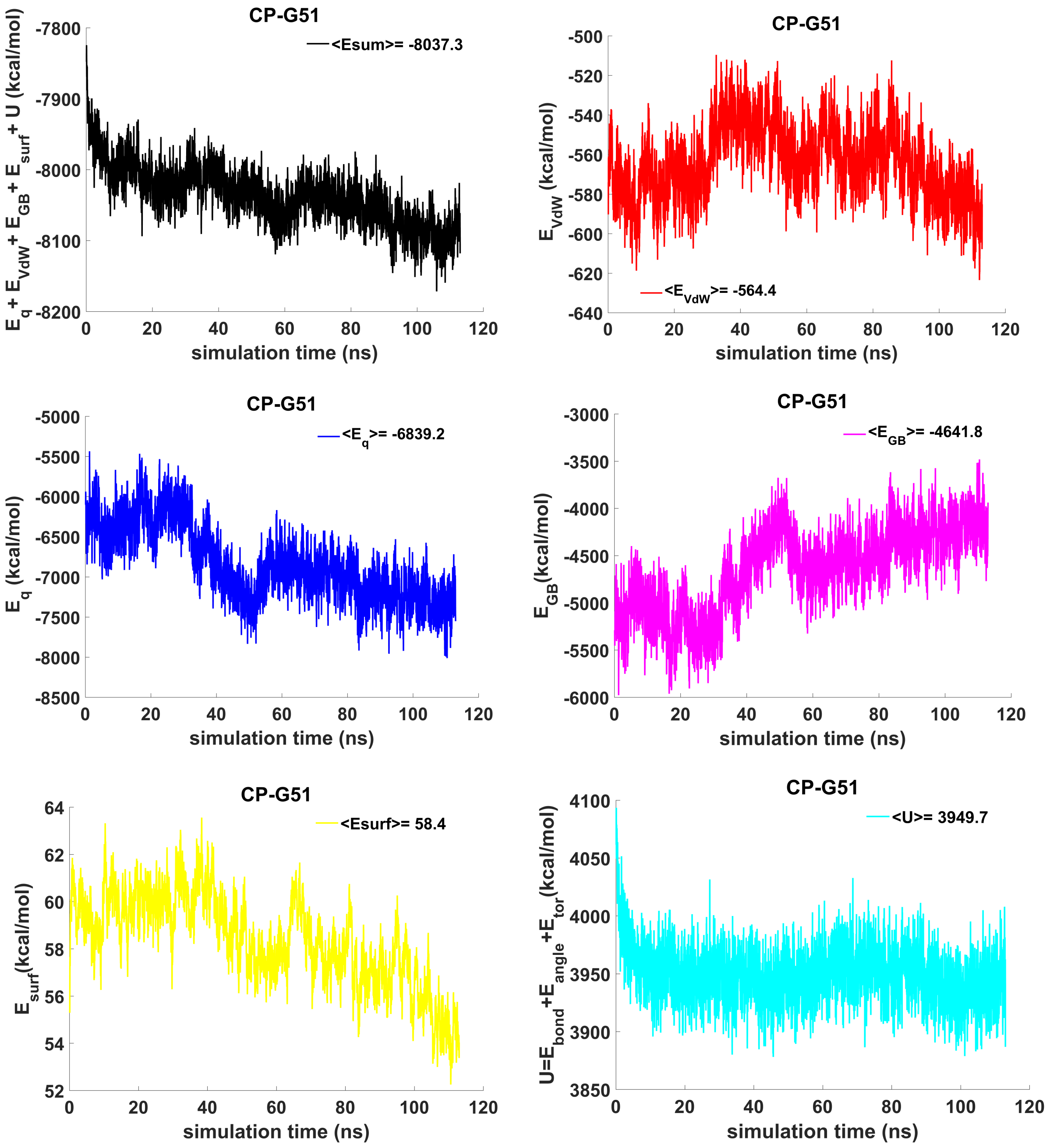


Fig.S1. B. **The contributions to free energy during MD calculations for the CP-G51 model**. $E_{eq}$ – electrostatic, $E_{vdw}$ - Van der Waals, $E_{GB}$ - polar energy of solvation, $E_{surf}$ - non-polar energy of solvation due to the hydrophobic surface available to the solvent, $U = E_{bond} + E_{angle} + E_{tor}$ , e.g. $E_{bond}$, $E_{angle}$ and $E_{tor}$ –bond, angle and torsion stress energies. The energy plots were smoothed using moving average method (span = 5). Average energy values are indicated in the figure legends.

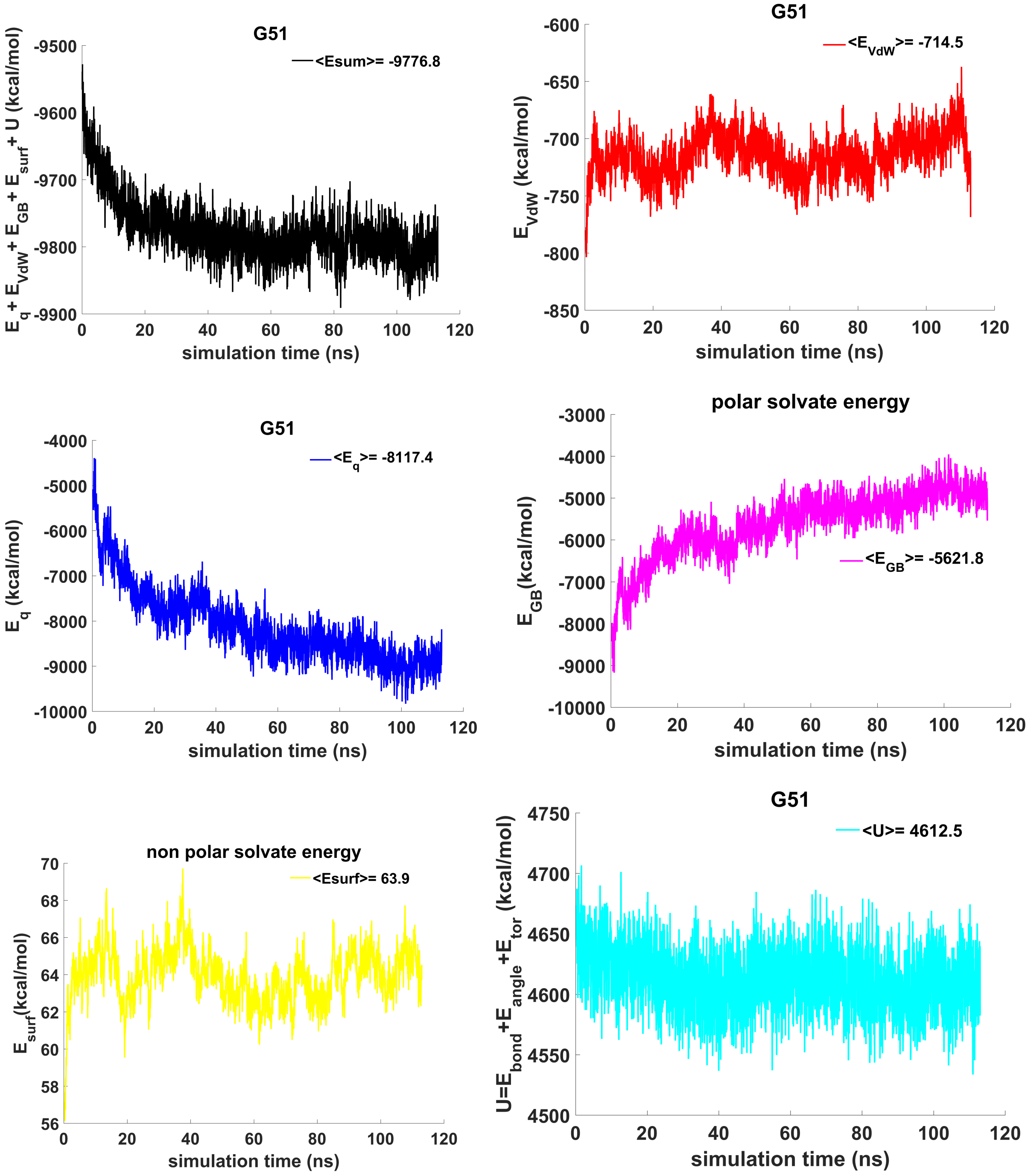


Fig.S1. C. **The contributions to free energy during MD calculations for the CP-G51 model**. $E_{eq}$ – electrostatic, $E_{vdw}$ - Van der Waals, $E_{GB}$ - polar energy of solvation, $E_{surf}$ - non-polar energy of solvation due to the hydrophobic surface available to the solvent, $U = E_{bond} + E_{angle} + E_{tor}$ , e.g. $E_{bond}$, $E_{angle}$ and $E_{tor}$ –bond, angle and torsion stress energies. The energy plots were smoothed using moving average method (span = 5). Average energy values are indicated in the figure legends.
